# Meta-analysis of sequence-based association studies across three cattle breeds reveals 25 QTL for fat and protein percentages in milk at nucleotide resolution

**DOI:** 10.1101/143404

**Authors:** Hubert Pausch, Reiner Emmerling, Birgit Gredler-Grandl, Ruedi Fries, Hans D. Daetwyler, Michael E Goddard

## Abstract

**Background:** Genotyping and whole-genome sequencing data have been collected in many cattle breeds. The compilation of large reference panels facilitates imputing sequence variant genotypes for animals that have been genotyped using dense genotyping arrays. Association studies with imputed sequence variant genotypes allow characterization of quantitative trait loci (QTL) at nucleotide resolution particularly when individuals from several breeds are included in the mapping populations.

**Results:** We imputed genotypes for more than 28 million sequence variants in 17,229 animals of the Braunvieh (BV), Fleckvieh (FV) and Holstein (HOL) cattle breeds in order to generate large mapping populations that are required to identify sequence variants underlying milk production traits. Within-breed association tests between imputed sequence variant genotypes and fat and protein percentages in milk uncovered between six and thirteen QTL (P<1e-8) per breed. Eight of the detected QTL were significant in more than one breed. We combined the association studies across three breeds using meta-analysis and identified 25 QTL including six that were not significant in the within-breed association studies. Closer inspection of the QTL revealed that two well-known causal missense mutations in the *ABCG2* (p.Y581S, rs43702337, P=4.3e-34) and *GHR* (p.F279Y, rs385640152, P=1.6e-74) genes were the top variants at two QTL on chromosomes 6 and 20. Another true causal missense mutation in the *DGAT1* gene (p.A232K, rs109326954, P=8.4e-1436) was the second top variant at a QTL on chromosome 14 but its allelic substitution effects were not consistent across three breeds analyzed. It turned out that the conflicting allelic substitution effects resulted from flaws in the imputed genotypes due to the use of a multi-breed reference population for genotype imputation.

**Conclusions:** Many QTL for milk production traits segregate across breeds. Metaanalysis of association studies across breeds has greater power to detect such QTL than within-breed association studies. True causal mutations can be readily detected among the most significantly associated variants at QTL when the accuracy of imputation is high. However, true causal mutations may show conflicting allelic substitution effects across breeds when the imputed sequence variant genotypes contain flaws. Validating the effect of known causal variants is highly recommended in order to assess the ability to detect true causal mutations in association studies with imputed sequence variant genotypes.

## Background

Whole-genome sequencing data have been generated for a large number of individuals from diverse cattle breeds. Many of the sequenced animals were selected in a way that they account for a large proportion of the genetic diversity of the entire population in order to ensure that the information content of the sequencing data is high [1]. This so-called “key-ancestor”-approach should reveal most polymorphic sites that segregate within breeds, at least the not too rare ones [2]. The availability of a comprehensive catalogue of sequence variants that segregate within breeds proved to be useful to pinpoint deleterious mutations particularly for recessive traits [3,4].

International consortia such as the 1000 bull genomes project (http://www.1000bullgenomes.com/) collected sequencing data from hundreds to thousands of individuals in order to characterize sequence variation that segregates within and across populations [5]. The fifth run of the 1000 bull genomes project provided genotypes at 39 million polymorphic sites for 1577 individuals that represent the most important dairy and beef breeds in the world. Reference panels that include sequence information from many breeds allow us to impute sequence variant genotypes at high accuracy for animals that have been genotyped using dense genotyping arrays [6,7]. Association studies with imputed sequence variant genotypes may facilitate to pinpoint causal mutations for complex traits [7,8].

Although sequence-based association studies uncovered many QTL (*e.g.*, [9–11]), little is known regarding their molecular-genetic underpinnings because the characterization of putatively causal variants was rarely attempted (*e.g.*, [5,12,13]). Sequence-based association studies typically reveal nearly identical P values for many adjacent variants that are in high linkage disequilibrium (LD). Such a pattern prevents differentiation between true causal mutations and anonymous markers that are in LD with them. Because many QTL for complex traits reside in non-coding regions of the genome, a functional prioritization of significantly associated sequence variants is not always possible [14].

Association studies that include animals from different breeds improve the resolution of QTL mapping because LD is conserved only over short distances across breeds [15]. However, trait definitions and data recording methods have to be standardized across breeds in order to allow for multi-breed association testing [16]. When restricted access to individual-level data precludes multi-breed association testing, meta-analysis enables us to combine summary statistics of association studies across populations thereby providing high power to detect QTL [17].

In this paper, we report on association studies between imputed sequence variant genotypes and two dairy traits in 17,229 animals from three cattle breeds. Meta analysis of association studies across three breeds allowed us to characterize 25 QTL for fat and protein percentages in milk at nucleotide resolution.

## Methods

### Animal ethics statement

No ethical approval was required for this study.

### Genotyped animals of the target populations

The target populations consisted of 1646 Braunvieh (BV), 6778 Fleckvieh (FV) and 8805 Holstein (HOL) bulls that had (partially imputed) genotypes at 573,650, 603,662 and 564,374 autosomal single nucleotide polymorphisms (SNPs). A subset of the animals (214 BV, 1475 FV and 345 HOL) was genotyped using the Illumina BovineHD (HD) bead chip that comprises 777,962 SNPs. All other animals were genotyped using the Illumina BovineSNP50 Bead chip (50K) that comprises 54,001 (version 1) or 54,609 (version 2) SNPs. The 50K genotypes were imputed to higher density using a combination of *Beagle* [18] and *Minimac* [19] (HOL, FV) or *FImpute* [20] (BV) as described previously [3,21]. To improve the accuracy of imputation, we increased the size of the reference panel by including HD genotypes of another 2070 FV and 842 BV animals that were available from previous projects. However, these data were not used for association analyses.

### Sequenced reference animals

Two reference panels were used to impute sequence variant genotypes for the animals of the target populations. Sequence variant genotypes for 8805 HOL animals were imputed using a multi-breed reference population that consisted of 1147 animals including 59 BV, 213 FV and 312 HOL animals that were available from the fourth run of the of the 1000 bull genomes project [5]. The sequencing reads were aligned to the UMD3.1 bovine reference genome using the *BWA-MEM* algorithm [22,23]. Single nucleotide polymorphisms, short insertions and deletions were genotyped for all sequenced animals simultaneously using a multi-sample variant calling approach that was implemented with the *mpileup* module of *SAMtools* [24] and that is described in Daetwyler et al. [5]. The sequence variant genotypes of 1147 reference animals were filtered to include 29,460,467 autosomal sequence variants with a minor allele frequency (MAF) greater than 0.0013 (*i.e.*, the minor allele was observed at least four times).

Sequence variant genotypes for 1646 BV and 6778 FV animals with (partially imputed) HD genotypes were imputed using a multi-breed reference population that consisted of 1577 animals that were included in the fifth run of the 1000 bull genomes project. The raw sequencing data were processed as described above. The sequence variant genotypes of 1577 reference animals were filtered to include 28,542,148 autosomal sequence variants that segregated in 123, 279 and 451 sequenced animals of the BV, FV and HOL breed, respectively.

A total of 24,180,002 sequence variants were a common subset of both reference populations. Sequence variant genotypes were imputed separately for each breed using a pre-phasing-based imputation approach that was implemented in the *Minimac* [25] software tool. Haplotype phases for the reference animals were estimated using *Beagle* (version 3.2.1) [18] (HOL) or *Eagle* (version 2.3) [26] (FV, BV). Sequence variants that were located between 71 and 78 Mb on chromosome 12 or between 23 and 30 Mb on chromosome 23 were not considered for association analyses because the accuracy of imputation was very low within both segments [7].

### Within-breed association testing

Association tests between imputed sequence variants and fat (FP) and protein percentages (PP) in milk were carried out for each breed separately using a variance components-based approach that was implemented in the *EMMAX* software tool and that accounts for population stratification and relatedness by fitting a genomic relationship matrix [27]. A genomic relationship matrix was built for each breed based on (partially imputed) HD genotypes using the method of Yang et al. [28] that was implemented in the *plink* (version 1.9) software tool [29]. Daughter yield deviations (DYDs) for FP and PP with an average reliability of 0.92 (±0.04) were the response variables in FV. Estimated breeding values (EBVs) with an average reliability of 0.89 (±0.12) and 0.95 (±0.03) were the response variables in BV and HOL. The phenotypic correlation between FP and PP was 0.53, 0.64 and 0.69 in BV, FV and HOL cattle. Predicted allele dosages were used as explanatory variables for the association tests. Sequence variants with P values less than 1e-8 were considered as significantly associated.

### Identification of QTL that segregate within and across breeds

Genomic regions with significantly associated variants were inspected manually. Genes that were annotated within 1 Mb intervals centered on the top variant were extracted from the UMD3.1 annotation of the bovine genome [30] using the Reference Sequence database (RefSeq release 82) from the National Center for Biotechnology Information (NCBI) and compared to known QTL for bovine milk production traits using literature review. Genomic regions were considered as across-breed QTL when significantly associated sequence variants (P<1e-8) were located within a 1 Mb interval centered on top variants that were detected at P<1e-8 in another breed.

### Meta-analysis of FP and PP across three breeds

Estimated allelic substitution effects and corresponding standard errors from the within-breed association studies (see above) were divided by phenotypic standard deviations to standardize the results of the association studies across breeds. Variants with an effect size greater than five standard deviations were not included in the metaanalysis (most of these variants also had low MAF and the large effect size might result from erroneously imputed alleles). The number of variants that were excluded because they had an effect size greater than five standard deviations varied across breeds and traits and ranged from 13,461 to 22,079. Meta-analysis was performed using an inverse variance-based approach that takes into account sample size, allelic substitution effect and standard error [17]. Heterogeneity of the effect sizes across breeds was evaluated using Cochran’s Q test [31] that was implemented in the *METAL* software package [17]. The functional consequence of significantly associated sequence variants was predicted using the *Variant Effect Predictor* tool from Ensembl [32]. Variant-specific estimates of FST were calculated for 25 QTL and whole genome sequence variants using 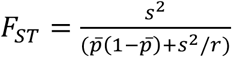, where s^2^ is the sample variance of allele frequency between breeds, 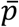 is the mean allele frequency across breeds and r is the number of breeds [33,34].

### Validation of 25 QTL in another population

A validation population that consisted of 1839 FV cows was genotyped at 777,962 SNPs using the HD bead chip. Haplotype phases were estimated using *Beagle* [26]. Sequence variant genotypes were imputed using *Minimac* [19] considering 1147 animals from the fourth run of the 1000 bull genomes project as a reference population (see above). EBVs for FP and PP with an average reliability of 0.50 (±0.03) were used as response variables for the association tests using the *EMMAX* [27] software tool as described above. Variants that had P values less than 0.05 and allelic substitution effects that were in the same direction as in the meta-analysis were considered to be validated in the cow population.

## Results

Genotypes for more than 28 million sequence variants were imputed for 17,229 progeny-tested bulls of the BV, FV and HOL cattle breeds using a population-based genotype imputation approach. Following genotype imputation, we considered 18,063,587, 19,021,606 and 17,318,499 imputed sequence variants that had MAF greater than 0.005 in BV, FV and HOL, respectively, for association testing. Between 32.4 and 35.1% of the imputed sequence variants had MAF less than 0.05 (see **Additional file 1 Figure S1**). The estimated mean (and median) accuracy of imputation (r^2^-values from *Minimac*) was 0.78 (0.96), 0.80 (0.97) and 0.79 (0.99) in BV, FV and HOL, respectively, and 85.4, 86.7 and 83.6% of the variants were imputed at an estimated accuracy greater than 0.3.

**Figure 1.**
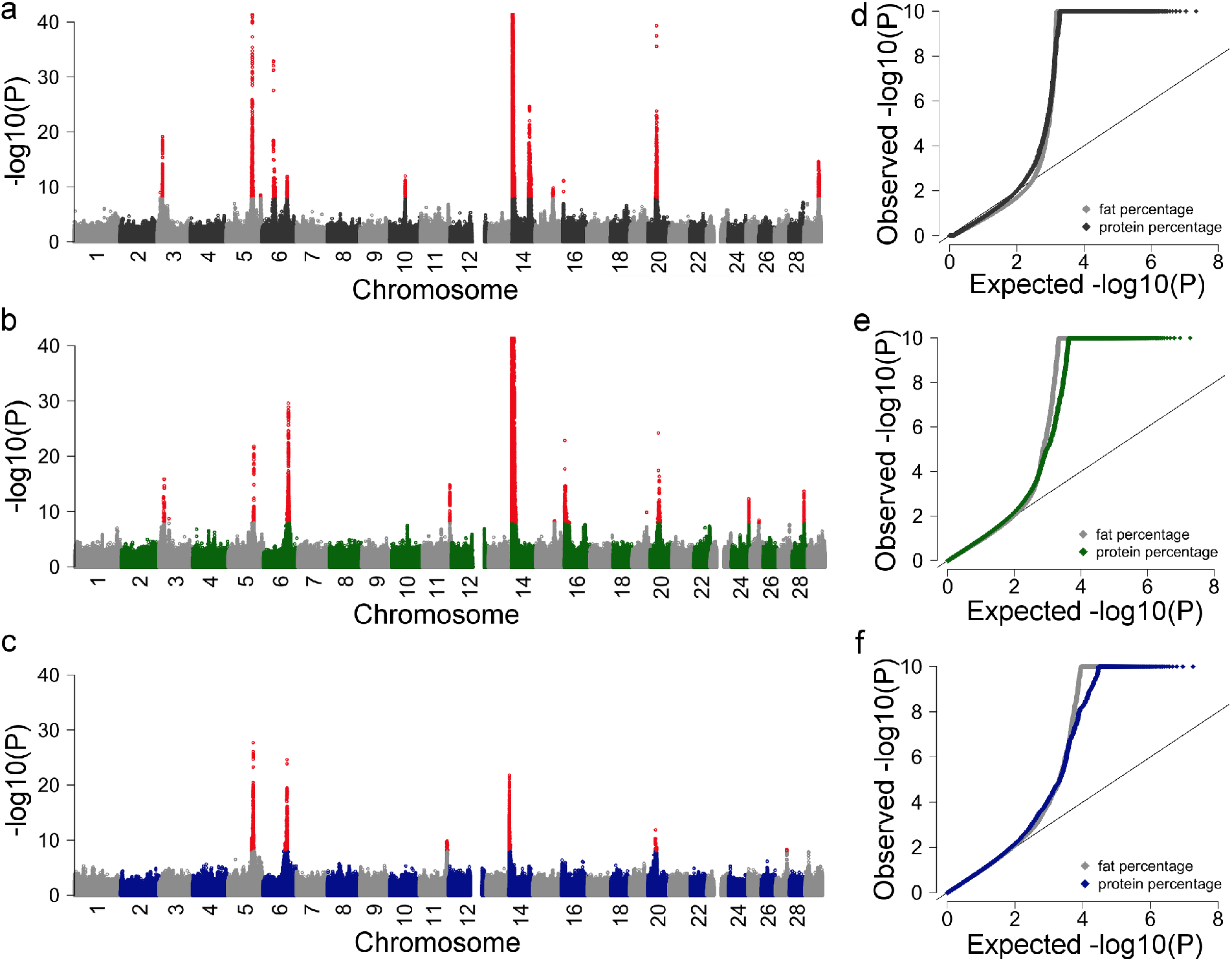
Detection of QTL for dairy traits in three cattle breeds. (a-f) Composite manhattan and corresponding quantile-quantile plots showing the association of 17,318,499, 19,021,606 and 18,063,587 imputed sequence variants, respectively, with FP and PP in HOL (a, d), FV (b, e) and BV (c, f). The composite manhattan plots summarize the results for FP and PP, *i.e.*, each dot shows the more significant P value that was observed across both traits. Red colours represent sequence variants with P values less than 1e-8. The y-axis is truncated at –log10(1e-40).

### Within-breed association studies for fat and protein percentages in milk

Association tests between imputed sequence variants and FP and PP were carried out separately for each breed. The number of associated sequence variants was higher in HOL and FV than BV which was likely because of a greater sample size in HOL and FV (BV: 1646, FV: 6778, HOL: 8805); 3249, 11,939 and 15,857 sequence variants were associated (P<1e-8) with FP in BV, FV and HOL, respectively, and 2296, 6515 and 15,674 were associated with PP. The difference in the number of significant variants per trait is mostly attributable to the properties of individual QTL regions. This is exemplified by the *DGAT1* QTL on BTA14, which has a more pronounced effect on FP than PP. In Fleckvieh and Holstein, respectively, 10,046 and 13,401 variants at the proximal region (<10 Mb) of chromosome 14 were associated (P<1e-8) with FP whereas only 2799 and 9493 variants were associated with PP. Two, 3106 and 10,029 variants were significant for both traits and 5543, 15,348 and 21,502 were significant for at least one trait in BV, FV and HOL, respectively.

Six, thirteen and twelve QTL, respectively, were detected in BV, FV and HOL cattle (Figure 1). Eleven QTL were detected in one breed, five and two QTL were detected in two and three breeds, respectively (see **Additional file 2 Table S1**). However, the top variants at QTL that were significant in more than one breed differed for all but one QTL; rs385640152 was the top variant for a QTL on chromosome 20 in all three breeds analysed.

Although more than 32% of the sequence variants had MAF between 0.005 and 0.05 (see above), only six (19%) QTL had MAF less than 0.05 (see Additional file 2 Table S1). QTL with MAF less than 0.05 were not detected in BV likely because the number of genotyped animals was too low to detect low-frequency QTL.

435 and 201 sequence variants, respectively, were significant (P<1e-8) for FP and PP in all three breeds analysed. Nineteen variants that were associated with FP in all breeds were located on chromosome 5 within the 5’-upstream sequence or intronic regions of *MGST1* (*microsomal glutathione S-transferase 1*). Another 415 variants that were significant for FP in all three breeds were located on chromosome 14 between 1,322,209 and 3,382,844 bp. In FV and HOL, several variants with P values less than 5e-306 (*i.e.*, the smallest possible P value that can be obtained using the *EMMAX* software tool) including the p.A232K-variant (rs109326954) in the *DGAT1* (*diacylglycerol O-acyltransferase 1*) gene were located within this segment. When variants with P values less than 5e-306 were ranked according to their t values (*i.e.*, regression coefficient divided by its standard error), rs109326954 was the second top variant in FV and its t-value (53.06) was only slightly less than the top variant (t=53.08, rs209876151 at 1,800,439 bp). In HOL, the t-value of rs109326954 was 2.7 points less than the top variant (rs110568020 at 1,699,681 bp). rs109326954 was not significant for FP in BV. At a QTL on chromosome 20, the p.F279Y-variant (rs385640152 at 31,909,478 bp) in the *GHR* (*growth hormone receptor*) gene was the top variant in all three breeds analysed.

221 non-coding sequence variants that were located between 87,154,594 and 87,434,710 bp on chromosome 6 and rs385640152 in the *GHR* gene were significant for PP in all breeds analysed.

Considering the allelic substitution effects of 18,063,587, 19,021,606 and 17,318,499 sequence variants in BV, FV and HOL, respectively, the correlation between FP and PP was 0.52, 0.61 and 0.53. The correlation between the FP and PP allelic substitution effects of 19 detected QTL was 0.81 (Figure 2). The largest effects were detected for QTL on chromosomes 14, 20 and 6 that encompassed the *DGAT1*, *GHR*, *ABCG2* (*ATP binding cassette subfamily G member 2*) and *CSN1S1* (*casein alpha s1*) genes. The QTL on BTA14 encompassing the *DGAT1* gene had larger effects on FP than PP whereas the effects of the QTL on BTA6 and BTA20 were more pronounced for PP than FP.

### Multi-breed meta-analysis uncovers six additional QTL

Meta-analysis of the within-breed association studies across three breeds revealed 16,086 and 14,020 sequence variants that were significantly associated (P<1e-8) with FP and PP, respectively (see **Additional file 3 Table S2 & Additional file 4 Table S3**). 23,786 variants were significant for at least one trait and 6320 variants were significant for both traits. The significant sequence variants clustered at 25 QTL including six that did not meet the significance threshold (P<1e-8) in any of the within-breed association studies (Table 1, Figure 3).

**Figure 2.**
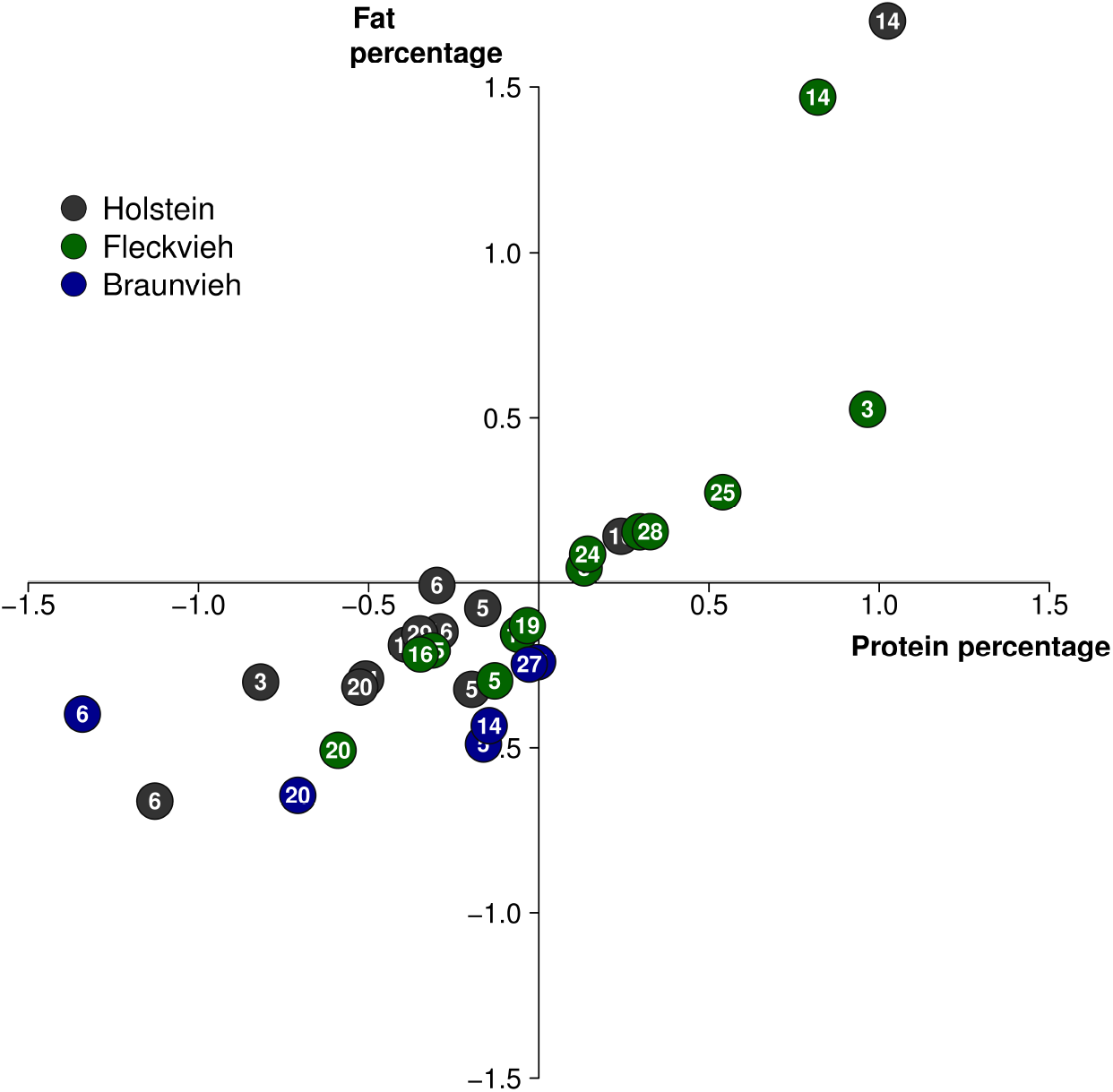
Effect of 31 QTL on fat and protein percentages in milk.

Allelic substitution effects of six, thirteen and twelve QTL, respectively, that were detected in HOL (grey), FV (green) and BV (blue) cattle. The allelic substitution effects of the top variants were divided by phenotypic standard deviations to standardize QTL effects across breeds.

The top variants at most QTL (20/25) had MAF greater than 0.005 in all three breeds analysed. However, the power of the within-breed association studies was likely not sufficient to detect all of them at P<1e-8 (see **Additional file 5 Table S4**). Seven QTL showed evidence of across-breed heterogeneity of the allelic substitution effects at P<0.05. The genetic differentiation of the three breeds was greater at 25 QTL (FST=0.079) than 24,180,002 genome-wide sequence variants (FST=0.059) indicating that at least some QTL were targets of recent selection.

The top variants at 19 QTL had MAF greater than 0.01 in a validation population that consisted of 1839 FV cows. The allelic substitution effects of 16 variants had P values less than 0.05 in the validation population and were in the same direction as in the meta-analysis (see **Additional file 5 Table S5**).

**Figure 3.**
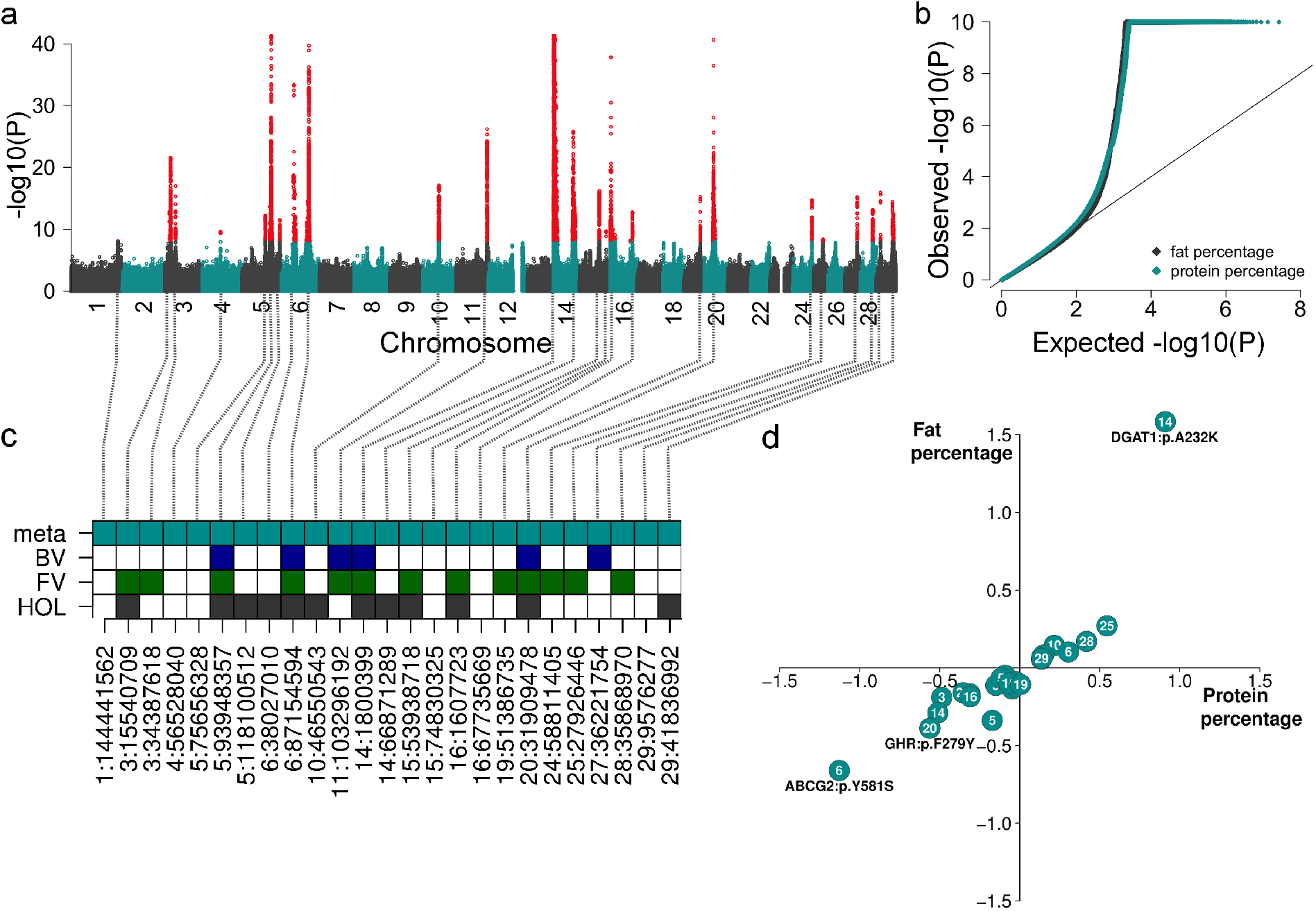
Meta-analysis of fat and protein percentages in milk across three cattle breeds. (a) Composite manhattan plot that shows the association of 26,473,121 imputed sequence variants with FP and PP in the meta-analysis. The composite manhattan plot summarizes the results of the meta-analyses, *i.e.*, each dot shows the more significant P value that was observed across both traits. Red colours represent sequence variants with P values less than 1e-8. The y-axis is truncated at –log10(1e-40). (b) Quantile-quantile plot of the meta-analyses. Grey and cyan colour represent P values of 26,473,121 imputed sequence variants for FP and PP, respectively. (c) Overview of 25 QTL that were significant at P<1e-8 in the meta-analysis and within-breed association studies. Filled squares indicate that QTL were significant in the respective analysis. The labels at the x-axis represent the positon of the top variant at each QTL. (d) Allelic substitution effects of 25 OTL on FP and PP. The QTL effects are given in phenotypic standard deviations. Bold type indicates three causal missense mutations in the *ABCG2*, *DGAT1* and *GHR* genes.

The most significantly associated variant at a QTL on chromosome 6 was a known causal mutation for milk production traits (p.Y581S, rs43702337 at 38,027,010 bp, P=4.3e-34) in the *ABCG2* gene [35]. The serine-encoding C-allele had a frequency of 0.008 in HOL and it decreased FP and PP (Figure 3d, Table 1). The serine variant did not segregate in the BV and FV bulls that had imputed sequence variant genotypes.

The p.F279Y-variant in the *GHR* gene (rs385640152 at 31,909,478, P=1.6e-74) was the top variant at a QTL on BTA20. Consistent with previous findings, the tyrosine encoding T-allele decreased FP and PP in all three breeds analysed [8,36]. The frequency of the tyrosine variant was 0.06, 0.05 and 0.15 in BV, FV and HOL (**see Additional file 2 Table S1**). The p.S18N variant (rs136247583) in the *PRLR* (*prolactin receptor*) gene [37] was polymorphic in the three breeds analysed and the frequency of the asparagine variant was 0.87 (HOL), 0.25 (FV) and 0.14 (BV). However, rs136247583 was neither associated with FP and PP in the meta-analysis (PFP=0.82, PPP=0.06) nor in any of the within-breed association studies (PFP>0.17, PPP>0.15).

A QTL on chromosome 1 was associated with FP in the meta-analysis but not in the within-breed association studies. Six imputed sequence variants that were located downstream of the *SLC37A1* (*solute carrier family 37 member 1*) gene had P values less than 1e-8. The variants associated with FP were in immediate vicinity to a QTL for phosphorus concentration in milk that was detected in an Australian Holstein cattle population [13]. The top variant (rs136426342 at 144,441,562 bp) of the metaanalysis was less than 75 kb away from two variants in high LD (rs109254133 at 144,367,474 bp and rs208161466 at 144,377,960 bp) that were plausible causal mutations in Holstein cattle [13]. However, their P values (P=3.2e-7 and P=8.0e-7) were higher than the P value (8.6e-9) of the top variant.

The meta-analysis revealed three distinct QTL on chromosome 5; 590 significantly associated sequence variants were located between 75,633,853 and 75,790,113 bp. This interval encompassed the *NCF4* (*neutrophil cytosolic factor 4*) and *CSF2RB* (*colony stimulating factor 2 receptor beta common subunit*) genes, which we did not consider as candidate genes for milk production traits. However, the top variant was less than 20 kb downstream of the translation end of the *TST* (*thiosulfate sulfurtransferase*) gene, which we considered as a functional candidate gene. Another QTL on BTA5 encompassed 1916 significantly associated sequence variants that were located between 88,235,301 and 98,233,638 bp. The effect of his QTL was more pronounced on FP than PP in all three breeds analysed. The most significantly associated variant (rs209372883 at 93,948,357 bp, P=9.8e-100) was located in the second intron of the *MGST1* gene. However, the top variant at that QTL differed across breeds. Another QTL on BTA5 encompassed 36 significantly associated sequence variants that were located between 118,086,877 and 118,264,313 bp including a missense mutation (p.R355C) in the *TBC1D22A* (*TBC1 domain family member 22A*) gene. However, the P value of the missense mutation was clearly higher (P=3.2e-9) than the most significantly associated non-coding variant (rs384440535 at 118,100,512 bp, P=3.0e-12).

The allelic substitution effects of most QTL were in the same direction for FP and PP (Table 1). However, two QTL on chromosomes 19 and 27 were highly significantly associated with FP (P<5.9e-16) but not with PP (P>0.1). The top variants at these QTL were located in intronic regions of the *FASN* (*fatty acid synthase*) and *AGPAT6* (*1-acylglycerol-3-phosphate O-acyltransferase 6*) genes.

### Conflicting effects of a known causal variant across breeds

The p.A232K-variant (rs109326954) in the *DGAT1* gene was the second top variant in the FP meta-analysis and its P value (8.4e-1436) was marginally higher than the top variant (rs208317364 at 1,800,399 bp, P=1.2e-1437). However, the allelic substitution effect of rs109326954 was not consistent across three breeds analysed. While the lysine variant increased FP and PP in FV and HOL, it decreased both traits in BV. The effect of rs109326954 on FP and PP was significant in FV and HOL but not in BV cattle (P>0.2). The lysine variant had a frequency of 0.08, 0.09 and 0.26 in the BV, FV and HOL animals, respectively, that had imputed genotypes. The allele frequency was similar in the sequenced FV (0.06) and HOL (0.30) animals. However, it was only 0.004 in 123 sequenced BV animals (*i.e.*, only one animal was heterozygous). Moreover, the accuracy of imputation (r^2^-value from *Minimac*: 0.02) was very low for rs109326954 in BV cattle. Taken together, these findings indicate that the imputed genotypes at rs109326954 were flawed in BV which caused the conflicting allelic substitution effects.

**Table 1:**
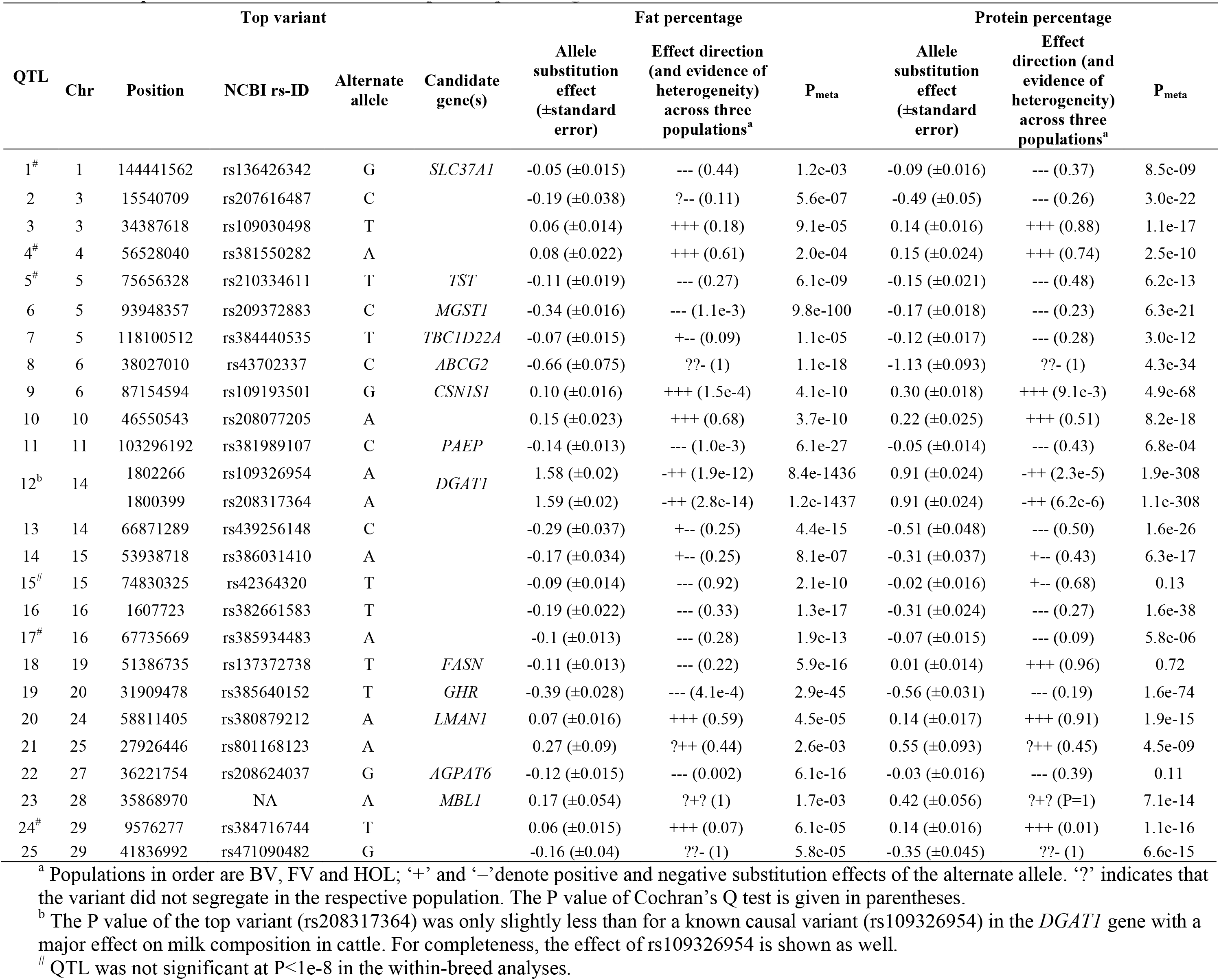
Top variants at 25 QTL for fat and protein percentages in milk.

## Discussion

Our meta-analysis of association studies for FP and PP across three cattle breeds identified 25 QTL including six that were not detected at P<1e-8 in the within-breed analyses. Our findings show that including data from different breeds can increase the power of association studies with imputed sequence variant genotypes which agrees with van den Berg et al. [15]. The power to detect QTL may be even greater in multibreed association studies [38]. However, access to individual-level genotype data was restricted in our study which prevented us from analyzing raw data from all three breeds simultaneously. Considering that our study included sequence variant genotypes of more than 17,000 animals, the identified QTL are likely to be the major genetic determinants of milk production traits in BV, FV and HOL cattle. We may have missed QTL with small effects because we applied a rather conservative significance threshold in the meta-analysis. Applying a less stringent threshold and QTL mapping approaches that consider all variants simultaneously may reveal QTL that explain only a small fraction of the genetic variation [39].

We identified a number of QTL for milk production traits that were previously detected in several cattle breeds, *e.g.*, QTL that were located nearby the *SLC37A1*, *MGST1*, *ABCG2*, *CSN1S1*, *PAEP* (*progestagen associated endometrial protein*), *DGAT1*, *FASN*, *GHR* and *AGPAT6* genes [5,7,8,12,13,21,35,40]. Using imputed sequence variant genotypes, Daetwyler et al. [5] identified a QTL for FP in early lactation in FV and HOL cattle that encompassed the *AGPAT6* gene. Our meta analysis also identified significantly associated sequence variants (including the variants reported by Daetwyler et al. [5]) in the 5’-upstream region of the *AGPAT6* gene corroborating that this region controls milk fat content in dairy cattle. However, our within-breed analyses did not detect that QTL in HOL and FV cattle at P<1e-8, although the animals of our study were from the same populations and our sample size was two and four times greater compared to Daetwyler et al. [5]. While Daetwyler et al. considered phenotypes for FP in early lactation as response variables, we used FP across the entire lactation. Milk fat content is under different genetic control across the lactation cycle [42] and the findings of Daetwyler et al. [5] and our study indicate that sequence variation nearby *AGPAT6* primarily controls fat content in early lactation which also agrees with the expression of *AGPAT6* at different stages of lactation [43].

Two putatively causal missense mutations in the *ABCG2* [35] and *GHR* [36] genes were the top variants at QTL on chromosomes 6 and 20, respectively, demonstrating that causal mutations can be readily identified in association studies with imputed sequence variant genotypes. Moreover, a well-characterized missense mutation in the *DGAT1* gene [44,45] was the second top variant at a QTL on chromosome 14 and its P value was only marginally higher than the top variant. Considering that the metaanalysis revealed three known causal mutations as the top (or second top) variants, it is likely that true causal mutations for milk production traits were among the significantly associated sequence variants. However, most associated variants resided in non-coding regions of the genome and a functional characterization of such variants was not attempted in our study. Nevertheless, including the trait-associated sequence variants of our meta-analysis in genomic predictions may improve the reliability of genomic predictions for dairy traits in cattle also for breeds other than BV, FV and HOL [46–48].

A well-characterized causal mutation (p.A232K, rs109326954) in the *DGAT1* gene [44,45] was the second top variant at a QTL on chromosome 14. In agreement with previous findings [8,44,45], the lysine variant was associated with higher FP and PP in FV and HOL. However, it had conflicting effects in BV cattle. A criterion for causality is consistency of allelic substitution effects across breeds. Strictly applying this criterion would lead to the exclusion of the p.A232K-variant as a plausible causal mutation. Closer inspection of the genotypes revealed that the lysine variant had a frequency of 0.004 and 0.08, respectively, in 123 sequenced and 1646 imputed BV animals. According to previous studies, the alanine variant is (nearly) fixed in BV cattle [45,49], which corroborates that the genotypes of the sequenced reference animals are correct. Since the p.A232K was imputed at very low accuracy (r^2^=0.02), it is likely that the conflicting allele frequencies and allelic substitution effects resulted from flaws in the imputed genotypes. Excluding variants with low r^2^-values would have removed the p.A232K variant from the BV population [50]. However, using too stringent cutoff values also carries the risk to exclude well-imputed genotypes [50,51]. Thus, we did not filter the imputed sequence variants based on their r^2^-values. We used reference populations that included animals from diverse populations to impute sequence variant genotypes. Multi-breed reference panels include many sequence variants that are not polymorphic in the target population. While multibreed reference panels enable us to impute genotypes at high accuracy (e.g.,[6,7,52]), our findings also show that they may promote the imputation of alleles that do not segregate in the target population. It might be advisable to compile breed-specific imputation reference panels that include animals from diverse breeds but only variants that segregate in the sequenced animals from the target breed [7,53]. Such an approach would likely remove from the mapping population a significant proportion of variants with flaws in the imputed genotypes [53]. However, it is only applicable when many individuals of the target breed have been sequenced in order to determine which sequence variants are polymorphic in that specific population.

Not all QTL identified in the meta-analysis (Table 1) were significant in all breeds. This could be because a QTL does not segregate in the three breeds analyzed or because we lacked power to show that it was significant at P<1e-8 particularly if it had a low frequency [54]. At five QTL, the top variant was not polymorphic in all breeds and so most likely the QTL does not segregate in those breeds. In addition, *DGAT1* had one very rare allele in BV which was likely to be imputed erroneously. Apart from these six QTL, there is only one case where the QTL does not have an effect in the same direction in all breeds for the more significant trait. This suggests that most detected QTL segregate in all three breeds even though they were not significant in the within-breed analysis.

The correlation between the effects of these QTL on FP and PP is a little surprising because the pathways for fat and protein synthesis in milk are quite different. One reason for this correlation is a QTL that affects milk volume without an equally large change in fat or protein yield and hence changes both PP and FP in the same direction. *ABCG2* and *GHR* might be examples of this [35,36]. Some QTL which have a functional role in fat synthesis (*DGAT1*, *AGPAT6*, *FASN*) have a bigger effect on FP than PP but still have an effect on PP except for *FASN*. Conversely, *CSN1S1* and *PAEP* are major milk protein components yet they affect both traits in our study. Perhaps the correlation between PP and FP is due in part to competition for substrate between lactose synthesis (which drives milk volume) and fat or protein synthesis so that an increase in either fat or protein can cause a decrease in volume as seen in the case of *DGAT1* [44,45].

## Conclusions

Many QTL for milk production traits segregate across cattle breeds and meta-analysis of association studies across breeds has greater power to detect such QTL than within-breed association studies. Sequence variants that are associated with dairy traits often reside in non-coding regions of the genome. True causal variants at milk production QTL can be readily identified in association studies with accurately imputed sequence variant genotypes. However, using reference panels that include animals from many breeds to impute sequence variant genotypes for GWAS populations may also promote the imputation of alleles that are actually not polymorphic in the target population. Such flaws in the imputed sequence variant genotypes can result in inconsistent allelic substitution effects of true causal mutations across breeds thereby compromising the differentiation between true causal mutations and neutral sequence variants.

## Declarations

### Ethics approval and consent to participate

DNA for genotyping and sequencing was prepared from semen samples of artificial insemination (AI) bulls that were collected by approved commercial AI stations as part of their regular breeding and reproduction measures in cattle industry. No ethical approval was required for the present study.

### Consent to publication

Not applicable

## Availability of data and material

All relevant data supporting the results and conclusions are included in the manuscript and its supporting files. The raw genotype and phenotype data of the animals were provided by breeding organizations (see acknowledgements section) following the execution of material transfer agreements under the condition of strict confidentiality. Contact details of representatives of the breeding organizations are available from the corresponding author (Hubert Pausch). Sequence variant genotypes in the form of vcf-files were provided by the 1000 bull genomes consortium (http://www.1000bullgenomes.com/).

**List of abbreviations**
BV: Braunvieh
FP: fat percentage
FV: Fleckvieh
HD: high density
HOL: Holstein
LD: linkage disequilibrium
MAF: minor allele frequency
PP: protein percentage
QTL: quantitative trait loci
SNP: single nucleotide polymorphism

## Authors’ contributions

HP and MEG designed the experiments, HP analyzed the data, RE, BGG and RF provided genotype and phenotype data, HDD provided computing support, HP wrote the manuscript. All authors read and approved the final manuscript.

## Acknowledgements

We thank Arbeitsgemeinschaft Süddeutscher Rinderzüchter und Besamungsorganisationen e.V. (ASR), Arbeitsgemeinschaft österreichischer Fleckviehzüchter (AGÖF), Förderverein Bioökonomieforschung e.V. (FBF), German Holstein Association (DHV), Confederación de Asociaciones de Frisona Española (CONAFE), ZuchtData EDV Dienstleistungen GmbH, Vereinigte Informationssysteme Tierhaltung (vit) w.V. Verden and Braunvieh Schweiz for providing genotype and phenotype data. We acknowledge the 1000 bull genomes consortium for providing sequence variant genotypes for 1577 animals.

## Additional Files

### Additional file 1 Figure S1

File format: tif

Title: Allele frequency distribution of imputed sequence variants

Description: Blue, green and grey, respectively, represent the proportion of imputed sequence variants in BV, FV and HOL for ten allele frequency classes.

### Additional file 2 Table S1

File format: xlsx

Title: **Most significantly associated variants at 19 QTL**.

Description: Chromosomal position and allelic substitution effect of the alternate allele of the most significantly associated variant at six, thirteen and twelve QTL, respectively, that were detected in BV, FV and HOL cattle. The allelic substitution effects were divided by phenotypic standard deviations. Coloured background indicates variants that were top variants within a 1 Mb region. Please note that the causal mutation in the *DGAT1* gene (Chr14:1802266) is also presented although it was not the most significantly associated variant in any breed.

### Additional file 3 Table S2

File format: csv

Title: **Sequence variants associated (P<1e-8) with fat percentage**

Description: Chromosomal positon an P values of 16,086 sequence variants that wer significantly associated with fat percentage in the meta-analysis across three breeds.

### Additional file 4 Table S3

File format: csv

Title: **Sequence variants associated (P<1e-8) with protein percentage**

Description: Chromosomal positon an P values of 14,020 sequence variants that wer significantly associated with fat percentage in the meta-analysis across three breeds.

### Additional file 5 Table S4

File format: xlsx

Title: **Characteristics of 25 QTL in three breeds analysed**.

Description: Frequency of 25 QTL in the BV, FV and HOL cattle breeds. FST indicates the variant-specific genetic differentiation across breeds. The P value of the top variant is given for the more significant trait.

### Additional file 6 Table S5

File format: xlsx

Title: **Association of the top variants at 25 QTL with FP and PP in 1839 FV cows**.

Description: Characteristics of the top variants at 25 QTL in a validation population of 1839 FV cows. Bold type and ‘NA’ indicates P values less than 0.05 and variants that were not polymorphic

## References

1. Goddard ME, Hayes BJ. Genomic selection based on dense genotypes inferred from sparse genotypes. Proc Adv Anim Breed Genet. 2009;18:26–9.

2. Jansen S, Aigner B, Pausch H, Wysocki M, Eck S, Benet-Pagès A, et al. Assessment of the genomic variation in a cattle population by re-sequencing of key animals at low to medium coverage. BMC Genomics. 2013;14:446.

3. Pausch H, Kölle S, Wurmser C, Schwarzenbacher H, Emmerling R, Jansen S, et al. A nonsense mutation in TMEM95 encoding a nondescript transmembrane protein causes idiopathic male subfertility in cattle. PLoS Genet. 2014;10:e1004044.

4. Pausch H, Schwarzenbacher H, Burgstaller J, Flisikowski K, Wurmser C, Jansen S, et al. Homozygous haplotype deficiency reveals deleterious mutations compromising reproductive and rearing success in cattle. BMC Genomics. 2015;16:312.

5. Daetwyler HD, Capitan A, Pausch H, Stothard P, van Binsbergen R, Brøndum RF, et al. Whole-genome sequencing of 234 bulls facilitates mapping of monogenic and complex traits in cattle. Nat Genet. 2014;46:858–65.

6. Brøndum RF, Guldbrandtsen B, Sahana G, Lund MS, Su G. Strategies for imputation to whole genome sequence using a single or multi-breed reference population in cattle. BMC Genomics. 2014;15:728.

7. Pausch H, MacLeod IM, Fries R, Emmerling R, Bowman PJ, Daetwyler HD, et al. Evaluation of the accuracy of imputed sequence variant genotypes and their utility for causal variant detection in cattle. Gen Sel Evol. 2017;49:24.

8. Pausch H, Wurmser C, Reinhardt F, Emmerling R, Fries R. Short communication: Validation of 4 candidate causative trait variants in 2 cattle breeds using targeted sequence imputation. J Dairy Sci. 2015;98:4162–7.

9. Höglund JK, Buitenhuis B, Guldbrandtsen B, Lund MS, Sahana G. Genome-wide association study for female fertility in Nordic Red cattle. BMC Genetics. 2015;16:110

10. Tenghe AMM, Bouwman AC, Berglund B, Strandberg E, de Koning DJ, Veerkamp RF. Genome-wide association study for endocrine fertility traits using single nucleotide polymorphism arrays and sequence variants in dairy cattle. J Dairy Sci. 2016;99:5470–85.

11. Pausch H, Emmerling R, Schwarzenbacher H, Fries R. A multi-trait meta-analysis with imputed sequence variants reveals twelve QTL for mammary gland morphology in Fleckvieh cattle. Genet Sel Evol. 2016;48:14.

12. Littlejohn MD, Tiplady K, Fink TA, Lehnert K, Lopdell T, Johnson T, et al. Sequence-based association analysis reveals an MGST1 eQTL with pleiotropic effects on bovine milk composition. Sci Rep. 2016;6:25376.

13. Kemper KE, Littlejohn MD, Lopdell T, Hayes BJ, Bennett LE, Williams RP, et al. Leveraging genetically simple traits to identify small-effect variants for complex phenotypes. BMC Genomics. 2016;17:858.

14. Andersson L, Archibald AL, Bottema CD, Brauning R, Burgess SC, Burt DW, et al. Coordinated international action to accelerate genome-to-phenome with FAANG, the Functional Annotation of Animal Genomes project. Genome Biol. 2015;16:57.

15. van den Berg I, Boichard D, Lund MS. Comparing power and precision of within-breed and multibreed genome-wide association studies of production traits using whole-genome sequence data for 5 French and Danish dairy cattle breeds. J Dairy Sci. 2016;99:8932–45.

16. Seidenspinner T, Bennewitz J, Reinhardt F, Thaller G. Need for sharp phenotypes in QTL detection for calving traits in dairy cattle. J Anim Breed Genet. 2009;126:455–62.

17. Willer CJ, Li Y, Abecasis GR. METAL: fast and efficient meta-analysis of genomewide association scans. Bioinformatics. 2010;26:2190–1.

18. Browning BL, Browning SR. A unified approach to genotype imputation and haplotype-phase inference for large data sets of trios and unrelated individuals. Am J Hum Genet. 2009;84:210–23.

19. Howie B, Fuchsberger C, Stephens M, Marchini J, Abecasis GR. Fast and accurate genotype imputation in genome-wide association studies through pre phasing. Nat Genet. 2012;44:955–9.

20. Sargolzaei M, Chesnais JP, Schenkel FS. A new approach for efficient genotype imputation using information from relatives. BMC Genomics. 2014;15:478.

21. Frischknecht M, Pausch H, Bapst B, Signer-Hasler H, Flury C, Garrick D, et al. Highly accurate sequence imputation enables precise QTL mapping in Brown Swiss cattle. submitted

22. Li H. Aligning sequence reads, clone sequences and assembly contigs with BWA-MEM. Preprint at: http://arxiv.org/abs/1303.3997. Accessed at 4 July 2016.

23. Zimin AV, Delcher AL, Florea L, Kelley DR, Schatz MC, Puiu D, et al. A whole-genome assembly of the domestic cow, Bos taurus. Genome Biol. 2009;10:R42.

24. Li H, Handsaker B, Wysoker A, Fennell T, Ruan J, Homer N, et al. The Sequence Alignment/Map format and SAMtools. Bioinformatics. 2009;25:2078–9.

25. Das S, Forer L, Schönherr S, Sidore C, Locke AE, Kwong A, et al. Next-generation genotype imputation service and methods. Nat Genet. 2016;48:1284–7.

26. Loh P-R, Danecek P, Palamara PF, Fuchsberger C, Reshef YA, Finucane HK, et al. Reference-based phasing using the Haplotype Reference Consortium panel. Nat Genet. 2016;48:1443–8.

27. Kang HM, Sul JH, Service SK, Zaitlen NA, Kong S, Freimer NB, et al. Variance component model to account for sample structure in genome-wide association studies. Nat Genet. 2010;42:348–54.

28. Yang J, Lee SH, Goddard ME, Visscher PM. GCTA: a tool for genome-wide complex trait analysis. Am J Hum Genet. 2011;88:76–82.

29. Chang CC, Chow CC, Tellier LC, Vattikuti S, Purcell SM, Lee JJ. Second-generation PLINK: rising to the challenge of larger and richer datasets. GigaScience. 2015;4:7.

30. Florea L, Souvorov A, Kalbfleisch TS, Salzberg SL. Genome assembly has a major impact on gene content: A comparison of annotation in two bos taurus assemblies. PLoS ONE. 2011;6:e21400.

31. Cochran WG. The combination of estimates from different experiments. Biometrics. 1954;10:101–29.

32. McLaren W, Gil L, Hunt SE, Riat HS, Ritchie GRS, Thormann A, et al. The Ensembl variant effect predictor. Genome Biol. 2016;17:122.

33. Weir BS, Cockerham CC. Estimating F-statistics for the analysis of population structure. Evolution. 1984;38:1358–70.

34. Beissinger TM, Hirsch CN, Vaillancourt B, Deshpande S, Barry K, Buell CR, et al. A genome-wide scan for evidence of selection in a maize population under long term artificial selection for ear number. Genetics. 2014;196:829–40.

35. Cohen-Zinder M, Seroussi E, Larkin DM, Loor JJ, Everts-van der Wind A, Lee J-H, et al. Identification of a missense mutation in the bovine ABCG2 gene with a major effect on the QTL on chromosome 6 affecting milk yield and composition in Holstein cattle. Genome Res. 2005;15:936–44.

36. Blott S, Kim J-J, Moisio S, Schmidt-Küntzel A, Cornet A, Berzi P, et al. Molecular dissection of a quantitative trait locus: a phenylalanine-to-tyrosine substitution in the transmembrane domain of the bovine growth hormone receptor is associated with a major effect on milk yield and composition. Genetics. 2003;163:253–66.

37. Viitala S, Szyda J, Blott S, Schulman N, Lidauer M, Mäki-Tanila A, et al. The role of the bovine growth hormone receptor and prolactin receptor genes in milk, fat and protein production in Finnish Ayrshire dairy cattle. Genetics. 2006;173:2151–2164.

38. Lin DY, Zeng D. Meta-analysis of genome-wide association studies: no efficiency gain in using individual participant data. Genet Epidemiol. 2010;34:60–6.

39. MacLeod IM, Bowman PJ, Vander Jagt CJ, Haile-Mariam M, Kemper KE, Chamberlain AJ, et al. Exploiting biological priors and sequence variants enhances QTL discovery and genomic prediction of complex traits. BMC Genomics. 2016;17:144.

40. Wang X, Wurmser C, Pausch H, Jung S, Reinhardt F, Tetens J, et al. Identification and dissection of four major QTL affecting milk fat content in the German Holstein-Friesian population. PLoS ONE. 2012;7:e40711.

41. Littlejohn MD, Tiplady K, Lopdell T, Law TA, Scott A, Harland C, et al. Expression variants of the lipogenic AGPAT6 gene affect diverse milk composition phenotypes in Bos taurus. PLoS ONE. 2014;9:e85757.

42. Strucken EM, Laurenson YCSM, Brockmann GA. Go with the flow-biology and genetics of the lactation cycle. Front Genet. 2015;6:118.

43. Bionaz M, Loor JJ. ACSL1, AGPAT6, FABP3, LPIN1, and SLC27A6 are the most abundant isoforms in bovine mammary tissue and their expression is affected by stage of lactation. J Nutr. 2008;138:1019–24.

44. Grisart B, Coppieters W, Farnir F, Karim L, Ford C, Berzi P, et al. Positional candidate cloning of a QTL in dairy cattle: identification of a missense mutation in the bovine DGAT1 gene with major effect on milk yield and composition. Genome Res. 2002;12:222–31.

45. Winter A, Krämer W, Werner FAO, Kollers S, Kata S, Durstewitz G, et al. Association of a lysine-232/alanine polymorphism in a bovine gene encoding acyl-CoA:diacylglycerol acyltransferase (DGAT1) with variation at a quantitative trait locus for milk fat content. Proc Natl Acad Sci USA. 2002;99:9300–5.

46. Brøndum RF, Su G, Janss L, Sahana G, Guldbrandtsen B, Boichard D, et al. Quantitative trait loci markers derived from whole genome sequence data increases the reliability of genomic prediction. J Dairy Sci. 2015;98:4107–16.

47. van den Berg I, Boichard D, Lund MS. Sequence variants selected from a multi breed GWAS can improve the reliability of genomic predictions in dairy cattle. Genet Sel Evol. 2016;48:83.

48. VanRaden PM, Tooker ME, O’Connell JR, Cole JB, Bickhart DM. Selecting sequence variants to improve genomic predictions for dairy cattle. Genet Sel Evol. 2017;49:32.

49. Scotti E, Fontanesi L, Schiavini F, La Mattina V, Bagnato A, Russo V. DGAT1 p.K232A polymorphism in dairy and dual purpose Italian cattle breeds. Ital J Anim Sci. 2010;9:e16

50. Li Y, Willer CJ, Ding J, Scheet P, Abecasis GR. MaCH: using sequence and genotype data to estimate haplotypes and unobserved genotypes. Genet Epidemiol. 2010;34:816–34.

51. Liu Q, Cirulli ET, Han Y, Yao S, Liu S, Zhu Q. Systematic assessment of imputation performance using the 1000 Genomes reference panels. Brief Bioinform. 2015;16:549–62.

52. Howie B, Marchini J, Stephens M. Genotype imputation with thousands of genomes. G3. 2011;1:457–70.

53. Hayes B, Bowman P, Daetwyler H, Goddard M. Why can we impute some rare sequence variants and not others? Proc Adv Anim Breed Genet. 2015;21:41–4.

54. Sham PC, Purcell SM. Statistical power and significance testing in large-scale genetic studies. Nat Rev Genet. 2014;15:335–46.

